# Fiber density and matrix stiffness modulate distinct cell migration modes in a 3D stroma mimetic composite hydrogel

**DOI:** 10.1101/2021.02.27.433190

**Authors:** Harrison L. Hiraki, Daniel L. Matera, William Y. Wang, Alexander A. Zarouk, Anna E. Argento, Johanna M. Buschhaus, Brock A. Humphries, Gary D. Luker, Brendon M. Baker

## Abstract

The peritumoral stroma is a complex 3D tissue that provides cells with myriad biophysical and biochemical cues. Histologic observations suggest that during metastatic spread of carcinomas, these cues influence transformed epithelial cells, prompting a diversity of migration modes spanning single cell and multicellular phenotypes. Purported consequences of these variations in tumor escape strategies include differential metastatic capability and therapy resistance. Therefore, understanding how cues from the peritumoral stromal microenvironment regulate migration mode phenotypes has prognostic and therapeutic value. Here, we utilize a synthetic stromal mimetic in which matrix fiber density and bulk hydrogel stiffness can be orthogonally tuned to investigate the contribution of these two key matrix attributes on MCF10A migration mode phenotypes, epithelial-mesenchymal transition (EMT), and invasive potential. We developed an automated computational image analysis framework to extract migratory phenotypes from fluorescent images and determine 3D migration metrics relevant to metastatic spread. Using this analysis, we find that matrix fiber density and bulk hydrogel stiffness distinctly contribute to a variety of MCF10A migration modes including amoeboid, single mesenchymal, multicellular clusters, and collective strands. Taking advantage of the tunability of this material platform, we identify a combination of physical and soluble cues that induces distinct heterogeneous migration modes originating from the same MCF10A spheroid and use this setting to examine a functional consequence of migration mode – apoptotic resistance. We find that cells migrating as part of collective strands are more resistant to staurosporine-induced apoptosis than either disconnected multicellular clusters or individual invading cells. Improved models of the peritumoral stromal microenvironment that help elucidate relationships between matrix attributes and cell migration mode can contribute to ongoing efforts to identify efficacious cancer therapeutics that address migration plasticity-based therapy resistances.

## INTRODUCTION

Given the highly mechanical nature of solid tumor progression, a recent focus has been placed on physical features of the stroma and their role in cancer cell dissemination^1^. As breast cancer progresses towards metastatic disease, the stroma undergoes marked remodeling including concurrent and interrelated increases in collagen fiber density and tissue stiffness^2,3^. These hallmark changes of the tumor stroma have been previously implicated in driving epithelial-mesenchymal transition (EMT) signaling, influencing invasive and migratory processes, and enabling cancer cell escape via contact guidance or durotactic gradients^4–6^. However, the individual contributions of these cues on EMT and cell migration mode have not been studied in a setting that reflects the 3D organization and structure of stroma that surrounds the primary tumor. This is in part due to limitations of existing approaches to studying cancer cell escape. Intravital imaging and *ex vivo* tissue models enable faithful recapitulation of the tumor microenvironment, but contain many uncharacterized factors that obfuscate causal relationships between specific microenvironmental cues and cell behavior. *In vitro* models utilizing purified biopolymers such as type I collagen recapitulate aspects of the chemical composition of native tissue, but typically lack tunable and orthogonal control over disease-relevant features of the native extracellular matrix (ECM)^7^. In contrast, synthetic hydrogels offer a higher degree of tunability over chemical and mechanical properties but typically lack the fibrous topography of stromal ECM. Recent work by our group and others have established a composite approach to incorporating fibrous architecture in otherwise amorphous bulk hydrogels^6,8–11^.

In this work, we utilize a synthetic hydrogel composite to model the fibrous peritumoral stroma and investigate how orthogonal changes in fiber density and bulk hydrogel stiffness individually influence EMT, migration dynamics, and heterogeneity in migratory phenotypes. Cell-adhesive dextran vinyl sulfone (DexVS) fiber segments were encapsulated at controlled densities within amorphous, photocrosslinkable gelatin methacrylate (GelMA) bulk hydrogels with stiffness tunable over a range spanning normal to desmoplastic breast tissue^12,13^. 3D outgrowth of MCF10A spheroids (a non-tumorigenic mammary epithelial cell line) was studied to determine whether perturbations to physical cues from the stroma could activate a non-transformed cell type into an invasive state. A custom automated image analysis framework was developed to quantitatively assess migration metrics and multicellularity. Herein, we report the effects of orthogonally increasing fiber density and bulk stiffness on cell migration phenotype and dynamics. We find that decreasing the stiffness of nonfibrous bulk GelMA hydrogel promotes single cell escape via amoeboid migration while the presence of more rigid cell-adhesive fibers enables strand-like collective invasion. Intermediate levels of bulk stiffness and fiber density engender a diversity of migration strategies, reflecting the intratumoral heterogeneity observed histologically in patient tissues. Finally, we leverage microenvironmental settings that elicit heterogeneous migration modes to assess differential apoptosis response as a function of migration mode and find that cell escape via collective strands confers enhanced resistance to apoptotic stimuli.

## MATERIALS AND METHODS

### Reagents

All reagents were purchased from Sigma Aldrich and used as received, unless otherwise stated.

#### Mouse xenograft implantation and intravital microscopy

The University of Michigan Institutional Animal Care and Use Committee approved all animal procedures (protocol 00006795). The animals used in this study received humane care in compliance with the principles of laboratory animal care formulated by the National Society for Medical Research and Guide for the Care and Use of Laboratory Animals prepared by the National Academy of Sciences and published by the NIH (Bethesda, MD; publication no NIH 85-23, revised 1996). We established orthotropic tumor xenografts as we have previously described^14^. Briefly, 2×10^5^ MDA-MB-231 cells stably expressing LifeAct-GFP were injected into the fourth inguinal mammary fat pads of 17-21 week-old female NSG mice (The Jackson Laboratory). Orthotopic tumors were intravitally imaged for collagen and LifeAct-GFP by an Olympus FVMPE-RS upright microscope with Spectra-Physics Insight DS+ laser, 25 × NIR corrected objective, 512 × 512 matrix, 15% laser power, 4 μs dwell time, and 2.5× electronic zoom. Collagen and GFP signals were distinguished by excitation wavelengths 880nm and 920nm, respectively, and band pass filters of 460-500nm and 495-540nm, respectively.

#### Synthesis of modified dextran and gelatin

<u>DexVS :</u>Dextran was functionalized with vinyl sulfone pendant groups using a previously described protocol^9^. Briefly, linear high molecular weight dextran (MW 86,000 Da; MP Biomedicals) was reacted with pure divinyl sulfone (Fisher) under basic conditions (pH 13.0). Functionalization was terminated through pH adjustment to 5.0 with hydrochloric acid. <u>GelMA:</u> Methacrylated gelatin was synthesized according to a previously described protocol^15^. Briefly, type A porcine skin gelatin was dissolved in PBS at 60 °C. Methacrylic anhydride was added dropwise and the reaction terminated after 1 hr by 5x dilution with warm PBS. All reaction products were dialyzed against milli-Q water for 3 days at 37 °C, with water changed twice daily. Purified products were then lyophilized for 3 days and reconstituted at 100 mg mL^-1^ in PBS.

#### DexVS fiber segment fabrication

DexVS was dissolved at 0.6 g ml^-1^ in a 1:1 mixture of milli-Q water and dimethylformamide. Lithium phenyl-2,4,6-trimethylbenzoylphosphinate (LAP) photoinitiator (6 v/v %) and methacrylated rhodamine (2.5 v/v %) (Polysciences, Inc.) were added to the solution to facilitate photoinitiated crosslinking and fluorescent visualization, respectively. This polymer solution was electrospun in a humidity-controlled glove box held at 21 °C and 30-35% relative humidity. Electrospinning was performed at 0.25 ml hr^-1^ flow rate, 7 cm gap distance, and -7.0 kV voltage onto a grounded copper collective surface. Fiber layers were collected on glass cover slides and primary crosslinked under ultraviolet light (100 mW cm^-2^) for 60 sec. Fiber mats were detached from cover slides into milli-Q water and broken into individual fiber segments. Fibers were purified through a series of centrifugation steps to remove uncrosslinked polymer and entangled clumps of fibers before resuspension in a buffer (1 N NaOH, 1 M HEPES, 1 mg mL^-1^ phenol red in milli-Q water) at 10 v/v %. Prior to encapsulation within bulk hydrogels, fibers were coupled with 2.0 mM RGD (CGRGDS; CPC Scientific) via Michael-type addition to enable eventual cell adhesion.

#### Cell lines and culture

Human mammary epithelial cells MCF10A (ATCC) were cultured in DMEM/F12 (1:1) supplemented with 5 v/v % horse serum (Fisher), 20 ng mL^-1^ rhEGF (Peprotech), 0.5 mg mL^-1^ hydrocortisone, 100 ng mL^-1^ cholera toxin, and 10 μg mL^-1^ insulin (Fisher). MCF10As were passaged at confluency at a 1:4 ratio and used for studies until passage 8. Cells were cultured at 37 °C and 5% CO_2_. To generate a LifeAct-GFP expressing line, cells were infected with 3^rd^ generation lentivirus transducing pLenti.PGK.LifeAct-GFP.W (a gift from Rusty Lansford, Addgene plasmid #51010). Lentivirus was produced in HEK-293T human embryonal kidney cells (ATCC) using calcium phosphate based transfection of viral packaging and transgene plasmids. <u>Spheroid formation :</u>MCF10As were detached with 0.25% trypsin-EDTA (Life Technologies), counted, and formed into 200 cell-sized spheroids overnight in inverse pyramidal PDMS microwells (AggreWell™, Stem Cell Technologies) treated with 0.5% Pluronic F-127 to prevent cell adhesion.

#### Hydrogel Formation

GelMA hydrogels were formed at 5 w/v % under ultraviolet light (5 mW cm^-2^) in the presence of LAP. LAP concentration was varied from 0.15-0.3 mg ml^-1^ to modulate gel stiffness. Fibrous hydrogels were created by mixing the stock solution of fiber segments in the hydrogel precursor prior to gelation. Hydrogel stiffness and fiber density were independently modulated through the concentration of LAP or fiber stock in the hydrogel precursor solution, respectively.

#### Mechanical Testing

To measure gel microscale tensile mechanical properties, atomic force microscopy nanoindentation in contact mode was performed using a Nanosurf FlexBio atomic force microscope. A HYDRA6V-200NG (AppNano) probe tip with spring constant of 0.0322 N/m affixed with an 8.12 μm diameter glass microsphere (Fisher) was used. Indentations were performed in three distinct regions with 30 points taken within each region over a 100 x 100 μm grid (6 x 5). Control and fiber gels were crosslinking on glass coverslips and subject to nanoindentation measurements. Force-displacement curves were fit to the Hertz model assuming a Poisson’s ratio of 0.5 to determine Young’s Modulus values.

#### Migration studies

Spheroids were harvested and centrifuged to remove residual single cells. Spheroids (6,000 per mL of gel) and fiber segments (variable density) were simultaneously photoencapsulated in GelMA. Studies were cultured in complete MCF10A media for 2 or 4 days, replenishing media every other day. For matrix metalloproteinase (MMP) inhibition studies, gels were cultured in complete MCF10A media containing marimastat (50 mM). For EMT studies, gels were cultured in complete MCF10A media containing rhTGF-β1 (Invitrogen). For apoptosis studies, gels were cultured in complete MCF10A media containing staurosporine (Santa Cruz Biotechnology) and left on a rocker plate at 0.33 Hz for 6 hrs to enhance diffusive transport.

#### Fluorescence, staining, and microscopy

Samples were fixed with 4% paraformaldehyde for 1 hr at room temperature. To visualize the actin cytoskeleton and nuclei, samples were stained simultaneously with phalloidin and DAPI (Fisher) for 1 hr at room temperature. For immunostaining, gels were additionally permeabilized in PBS containing Triton X-100 (5 v/v%), sucrose (10 w/v%), and magnesium chloride (0.6 %w/v) and blocked in 4% BSA. Gels were then incubated in mouse anti-E-cadherin (1:500, Abcam #ab1416), rabbit anti-SNAI1 (1:1000, Cell Signaling Technologies #3879S), mouse anti-vimentin (1:500, Sigma #V63890), or rabbit anti-caspase-3 (1:500, ThermoFisher #700182) followed by Alexa-conjugated anti-mouse or anti-rabbit secondary antibodies for 8 hours each at room temperature. Fluorescent imaging was performed with a Zeiss LSM 800 laser scanning confocal microscope. For migration analysis, Z-stacks were acquired with a 10x objective. High-resolution images were acquired with a 40x objective. All images are presented as maximum intensity projections.

#### Cell migration analysis

Max intensity projections of spheroid nuclei and F-actin channels were inputs to a custom MATLAB code (Fig. 2A) which separately thresholded and object size filtered each channel to remove background (Fig. 2B). A user-drawn ellipsoidal ROI covering the spheroid body was used to separate the spheroid body from migratory cells within outgrowths. The user then confirmed each segmented F-actin structures to be an outgrowth, after which the code defined an outgrowth as contiguous or noncontiguous based on contiguity with the spheroid body (Fig. 2C). A separate function segmented overlapping nuclei to identify all nuclei within outgrowths (Fig. 2D). Individual outgrowth F-actin masks were used to determine migration distance into the surrounding hydrogel utilizing a separate custom function. Corresponding individual nuclei masks were used to determine nuclear counts and mark noncontiguous outgrowths as either multicellular clusters or single cells (Fig. 2E). All individual outgrowth nuclei and F-actin masks were then summed to produce final images of nuclei and F-actin channels (Fig. 2F-G). Individual outgrowth nuclei and F-actin masks were saved with counted nuclei or plotted lengths, respectively, and assigned an index to address discrepancies or outliers within final quantified data. Resulting data were stratified by migration mode and exported to a spreadsheet containing individual outgrowth indices, number of migratory cells, outgrowth areas, and migration distances (Fig. 2H). Finally, spheroid body and outgrowth masks were summed across all analyzed spheroids to produce heatmap overlays (Fig. 2I).

#### Statistics

Statistical significance was determined by one-way analysis of variance (ANOVA) with post-hoc analysis (Tukey test), with significance indicated by p < 0.05. All data are presented as mean ± standard deviation.

## RESULTS

### Fabrication of a tumor stroma mimetic with orthogonal control over key physical properties

Intravital imaging was used to characterize the fibrous architecture of the peritumoral stroma in a murine orthotopic xenograft tumor model^14^, revealing dense collagen fiber networks at the tumor-stroma interface (Fig. 1A). To model fibers of this length scale *in vitro*, DexVS was electrospun and processed into suspended fiber segments of comparable diameter^9^. DexVS fiber segments were encapsulated in GelMA bulk hydrogels to create a fibrous stroma mimetic surrounding embedded MCF10A spheroids (Fig. 1B), with orthogonal control over fiber density and bulk stiffness. The density of fibers within composite hydrogels was varied through the input volume fraction of fiber segments (Fig. 1C), while bulk hydrogel stiffness was controlled by the concentration of LAP photoinitiator present during polymerization. AFM nanoindentation confirmed that modulating fiber density did not affect composite gel stiffness (Fig. 1D) and that composites could be tuned over a range of 1.75 – 6.0 kPa, spanning values reported for healthy and metastatic breast tissue^12,13^ (Fig. 1E). This fiber-reinforced composite hydrogel system proved amenable to robust 3D cell migration in which cells engaged cell-adhesive fibers to invade the surrounding hydrogel (Fig. 1F).

**Fig.1.**
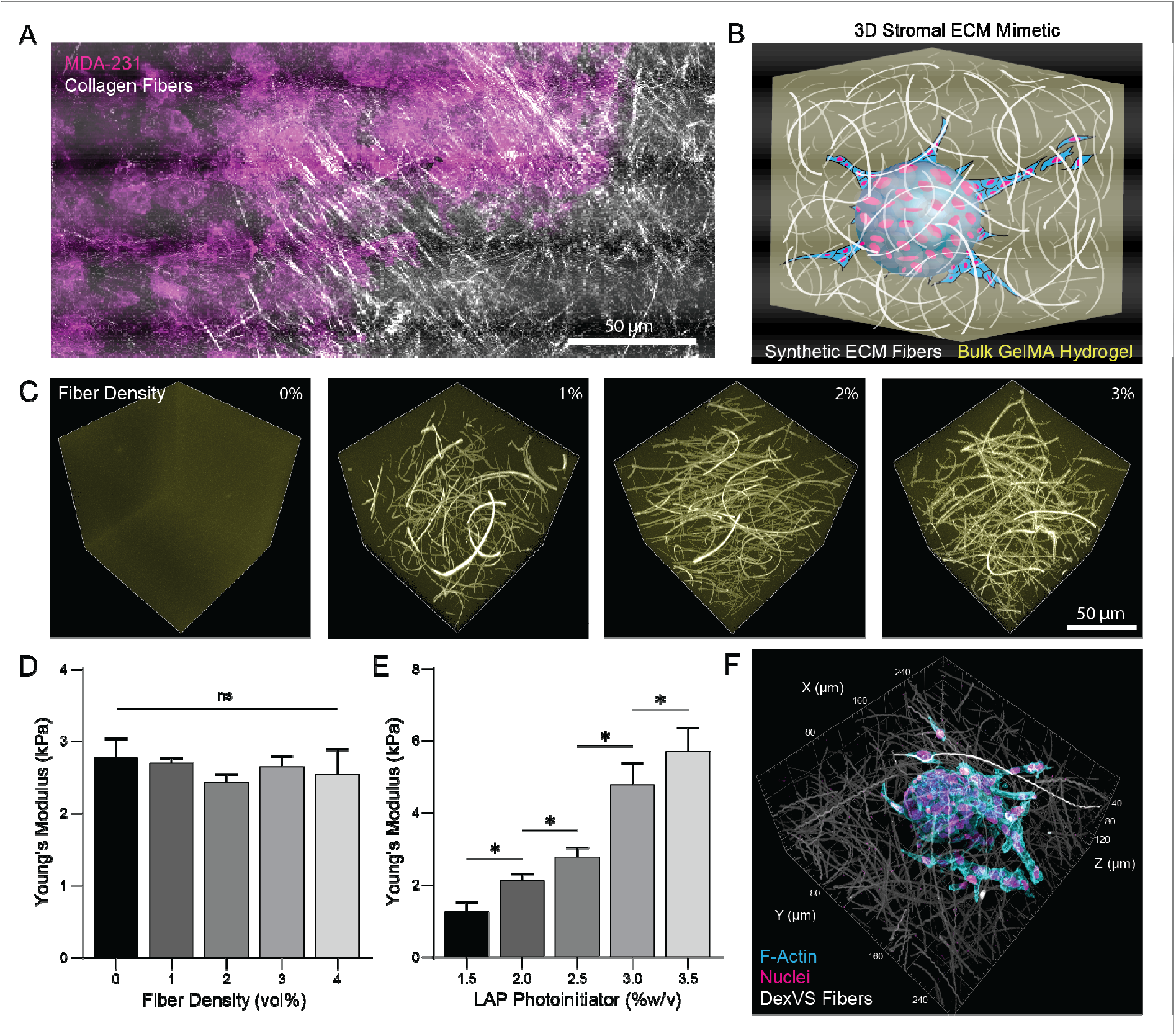
Fiber-reinforced hydrogel composites to model the peritumoral stroma. (A) Intravital image of peritumoral collagen fibers within a murine xenograft model. (B) Schematic diagram of spheroid outgrowth in stroma mimetic hydrogel composite. (C) 3D projections of confocal images of DexVS fiber densities within fluorescently labeled GelMA. 200 × 200 × 200 µm gel volume. Young’s modulus of hydrogels (D) at constant LAP concentration (2.5 w/v%) over a range of fiber densities or (E) over a range of LAP photoinitiator concentrations at constant fiber density (3 vol%) quantified via AFM nanoindentation. (F) Fluorescent image of MCF10A spheroid outgrowth in stroma mimetic.

**Fig.2.**
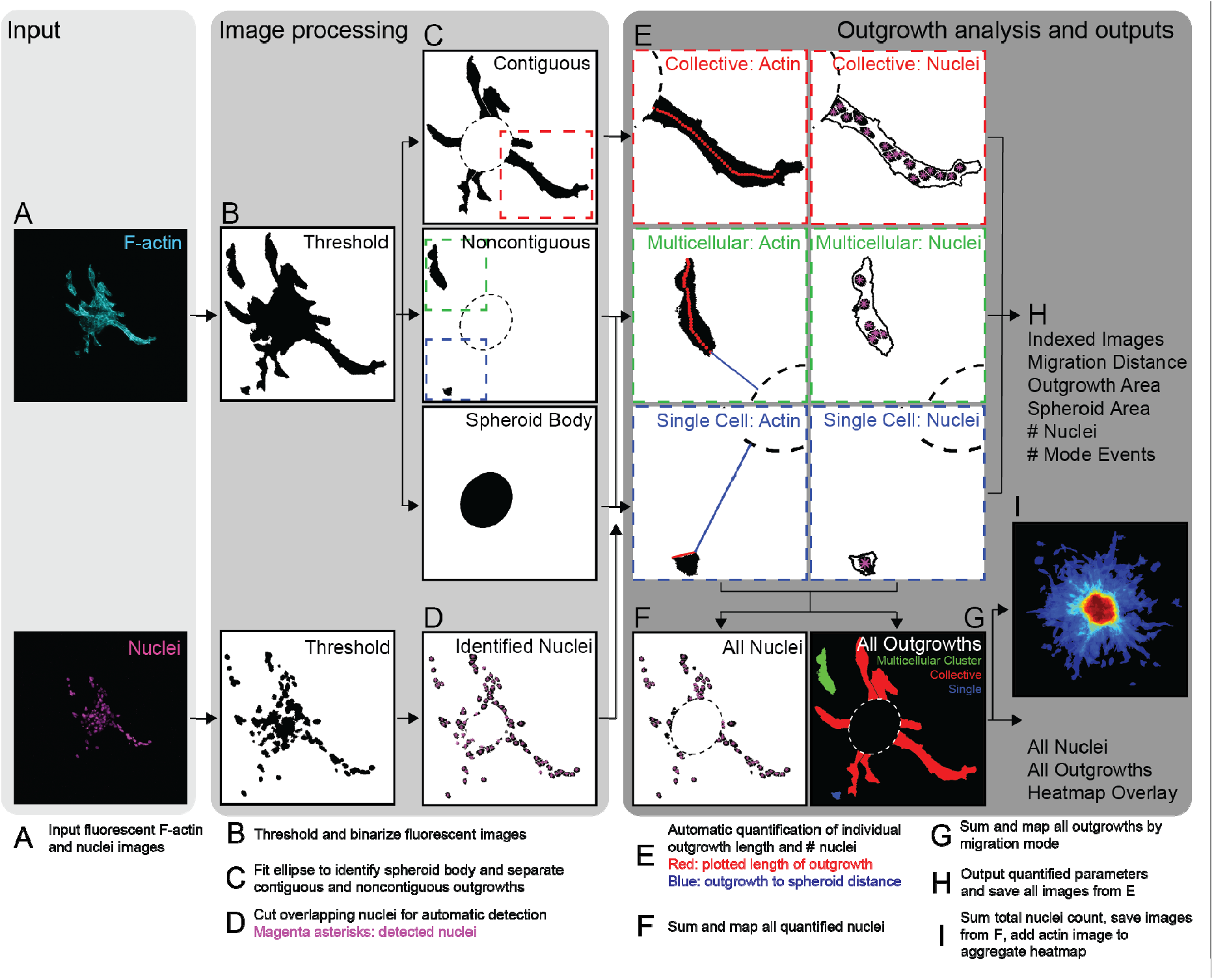
Flowchart of custom MATLAB code used to quantify spheroid outgrowth. (A) Code input requires fluorescent images of F-actin and nuclei. (B) Images are first subject to manual thresholding. A user-drawn ellipse is used to (C) segregate contiguous and noncontiguous outgrowths in the actin image and (D) identify all nuclei contained in outgrowths in the nuclei image. (E) Each outgrowth is automatically stratified by migration mode (collective, multicellular cluster, or single cell). Outgrowths are automatically analyzed individually to quantify area, invasion depth, and number of nuclei per outgrowth. Individual outgrowth actin and nuclei images are aggregated to output quantifications of (F) total number of migratory cells and (G) migration mode populations. Code outputs (H) a spreadsheet containing outgrowth quantifications as individual data points and spheroid averages; corresponding individual outgrowth images with plotted lengths and counted nuclei to check against quantified values; (I) generated images of heatmap overlays, all nuclei counted, and all outgrowths registered.

### Automated analysis of cell migration phenotype and invasion metrics relevant to cancer progression

To quantitatively assess migration metrics relevant to metastatic spread and the multicellularity or connectivity of cellular structures emanating from spheroids, we created an image analysis framework in MATLAB. The code processes F-actin- and nuclear-stained maximum projection fluorescent images (Fig. 2A) and requires minimal user image processing. Thresholded cellular and nuclear images (Fig. 2B) are segmented into distinct outgrowth structures and stratified by multicellularity (Fig. 2C-D). Invasion distances, outgrowth areas, and the number of nuclei within each distinct invading cellular structure are automatically quantified (Fig. 2E-G). Morphometrics and indexed images of invasive outgrowths with overlaid visualization of quantified invasion distance and nuclear count are then exported to address discrepancies in the data (Fig. 2H). If multiple spheroids are processed simultaneously, summed images of thresholded F-actin area are used to generate heatmap overlays for visualization of aggregate migration phenotypes within a single condition (Fig 2I). An in-depth explanation of the code can be found in the methods section.

Three key metrics were determined from raw images as quantitative measures of invasive potential and migration phenotypes: 1) total number of escaped cells, representing metastatic burden, as over a prolonged period of time all migratory cells can reach nearby vasculature to metastasize; 2) total migration distance, the summed migration distance of each cell from the spheroid periphery as a measure of net transtromal migration; and 3) migration mode percentage, providing a population distribution of migration modes emanating from individual spheroids. These metrics are used here to quantitatively describe cell migration dynamics following perturbations to mechanical and soluble cues within stromal ECM mimetics.

### Nonfibrous bulk hydrogels restrict migration to an amoeboid phenotype sensitive to bulk gel stiffness

As one of the earliest signs of breast cancer is tissue stiffening, we first investigated whether an increase in hydrogel stiffness/crosslinking, in the absence of fibrous architecture, was sufficient to induce invasive behavior of normally quiescent MCF10A spheroids^16,17^. Nonfibrous GelMA hydrogels spanning the range of Young’s moduli previously reported for healthy and metastatic states elicited solely single cell migration from MCF10A spheroids^12,13^ (Fig. 3A-B). Single migratory cells appeared rounded and frequently possessed a polarized F-actin-rich leading edge (Fig. 3C, white arrowheads). Timelapse imaging of cells stably expressing LifeAct-GFP to visualize actin cytoskeleton dynamics revealed cyclic blebbing and rapid translocations of cells, characteristic of amoeboid cell migration^18,19^ (Fig. 3D & Supp. Video 1). Migration dynamics were sensitive to hydrogel stiffness where increased stiffness hindered migration, evident by decreases in the number of cells migrating per spheroid and a reduction in total migration distance (Fig. 3E-F). Previous studies suggest amoeboid migration involves cells squeezing through matrix pores and therefore does not require matrix proteolysis^20^. To determine whether amoeboid migration within this biomaterial is similarly proteolysis independent, marimastat (MMS, a broad-spectrum MMP inhibitor) was added to culture media prior to the onset of migration. Although we noted a reduction in the number and invasion depth of escaped cells, cell escape was still observed with MMP inhibition (Fig. 3G). While stiffness/crosslinking of a cell-degradable, nonfibrous hydrogel clearly influenced the propensity for amoeboid-like single cell dissemination, there was surprisingly no evidence for multicellular migration that has been commonly observed histologically in carcinoma^21^.

**Fig.3.**
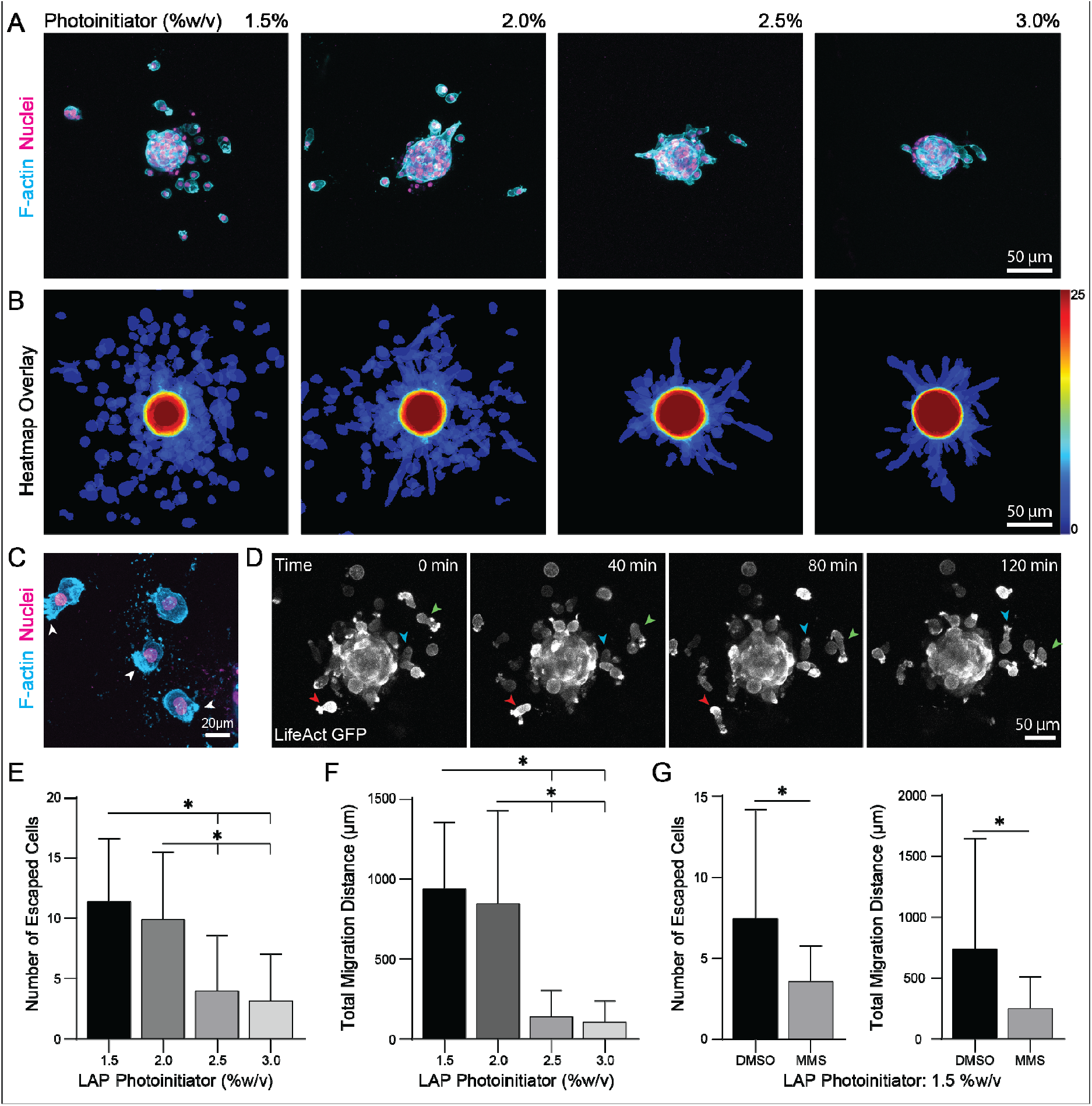
Nonfibrous, amorphous GelMA hydrogels favor single cell amoeboid migration with decreases in hydrogel stiffness/crosslinking. (A) Fluorescent images of spheroid outgrowth after 2 days in matrices crosslinked with different LAP photoinitiator concentrations. (B) Corresponding heatmap overlays created by an aggregate sum of binarized actin channels for n = 25 spheroids per condition. (C) Fluorescent image of single cells migrating with an amoeboid phenotype. White arrowheads indicate F-actin rich leading edges. (D) Timelapse of single cell migration within a 1.5% LAP (E ∼ 1.75 kPa) gel over a 2 hr range. Colored arrowheads track individual cells displaying cyclic blebbing migration. Quantification of (E) total number of escaped cells per spheroid and (F) total migration distance per spheroid measured as the sum of net migration distance of all nuclei. (G) Quantification of spheroid outgrowth in 1.5% LAP gels treated with broad-spectrum MMP-inhibitor marimastat (MMS). All data presented as mean ± std; * indicates a statistically significant comparison with p < 0.05.

### Fibrous architecture promotes collective strand migration

Fibrous architecture in the stroma undergoes substantial remodeling prior to and during metastasis highlighted by an increase in local fiber density at the tumor-stroma interface^2^. To determine if the presence of matrix fibers alone could initiate migration, we next modulated the density of DexVS fiber segments within highly crosslinked bulk GelMA hydrogels (E = 2.75 kPa), which limited migration in our previous study (Fig. 3). Interestingly, the inclusion of fibers over a range of densities (1-3 vol%) promoted MCF10A migration in the form of collective, multicellular strands (Fig. 4A-B), which engaged, recruited, and tracked along matrix fibers (Fig. 4C, red arrowheads). Increases in fiber density up to 3 vol% resulted in step-wise increases in the number of migrating cells per spheroid and total migration distance (Fig. 4D-E). At the two higher densities of matrix fibers tested (2 and 3 vol%), we noted a minor but significant increase in the number of outgrowths as single cells or multicellular clusters, here defined as any multinucleated structures that were noncontiguous with the spheroid (Fig. 4F). To confirm that direct integrin engagement of matrix fibers promoted migration, identical studies were repeated with hydrogels containing an equivalent 3 vol% density of fibers but lacking RGD functionalization (-RGD). Despite the availability of endogenous RGD throughout the bulk GelMA hydrogel, the absence of RGD presented on matrix fibers led to a significantly decreased frequency and total migration distance of collective strands (Fig. 4D-E). Taken together, these observations suggest that integrin engagement to stiff and mechanically anisotropic fibers is a requirement for coordinated multicellular invasion strategies as collective strands that remain connected to the spheroid or as disconnected multicellular clusters.

**Fig.4.**
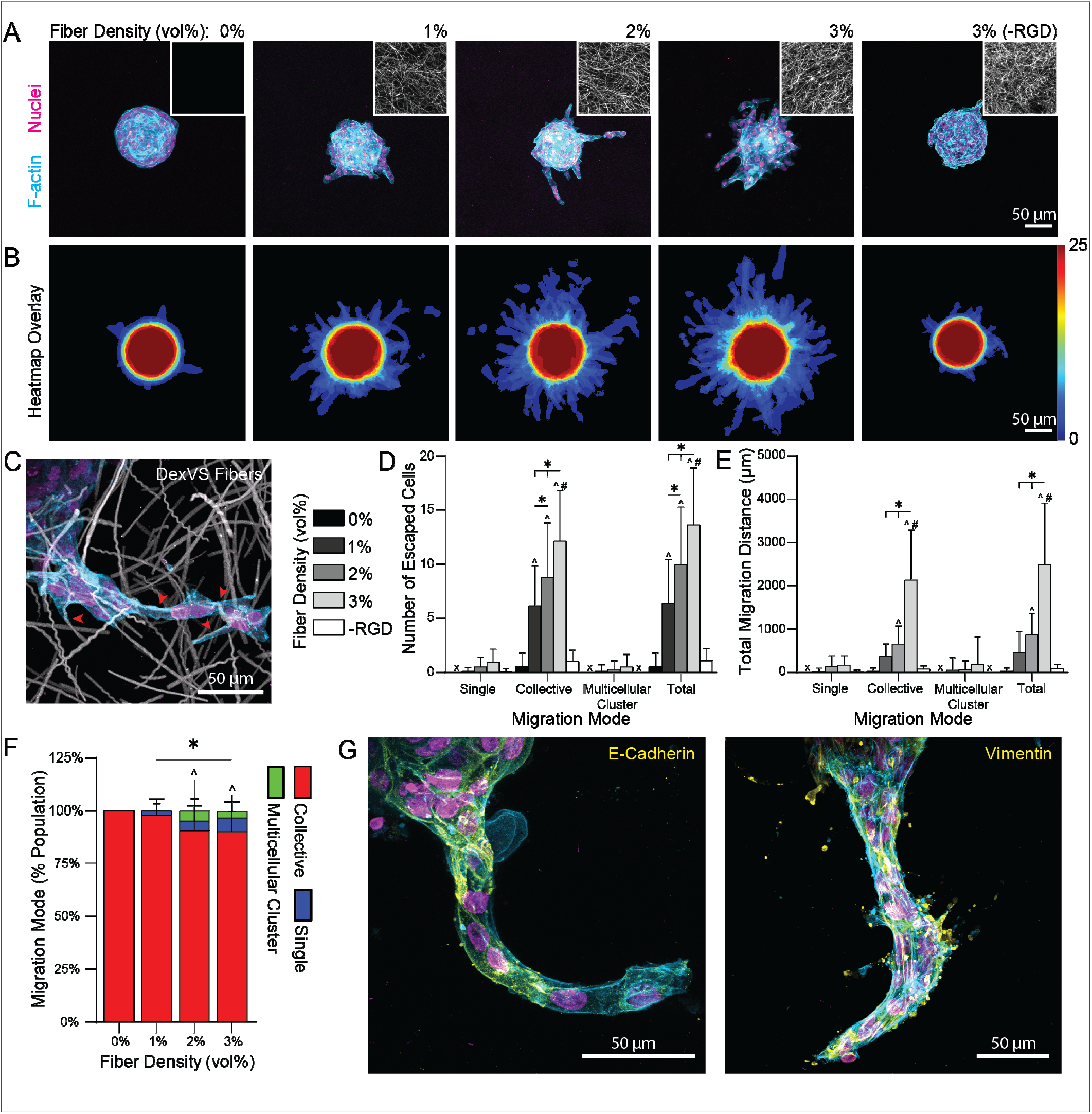
An increasing density of synthetic DexVS matrix fibers within amorphous GelMA promotes collective migration. (A) Fluorescent images of spheroid outgrowth over 4 days in stiff 2.5% LAP gels over a range of DexVS fiber densities. Insets: Representative 50 μm max projection of rhodamine-labelled DexVS fibers within each gel condition. (B) Corresponding heatmap overlays of binarized actin images for n = 25 spheroids per condition. (C) Representative fluorescent image of cells utilizing DexVS fibers functionalized with RGD to migrate from a spheroid. Red arrowheads indicate fiber engagement. Quantification of (D) total number of escaped cells and (E) total migration distance per spheroid as a function of migration mode. –RGD condition indicates 3 vol% fiber density gels containing fibers lacking RGD coupling. (F) Stratification of migration mode subpopulations per spheroid. (G) Representative images of multicellular outgrowths immunostained for E-cadherin and vimentin. All data presented as mean ± std; * indicates a statistically significant comparison with p < 0.05; ^ indicates significance against 0 vol% fiber density; # indicates significance against –RGD condition.

As mesenchymal cell migration of epithelial cells is largely believed to commence only after EMT, we immunostained collective strands for established epithelial (E-cadherin) and mesenchymal (vimentin) markers. Co-expression of both E-cadherin and vimentin suggested partial EMT in the formation and movement of these collective strands (Fig. 4G), compared to nonmigratory cells on the spheroid surface which only expressed E-cadherin (Supp. Fig. 1).

### Fibrous architecture and bulk stiffness regulate amoeboid vs. collective strand-like migration

As the presence of fibers within stiffer, more crosslinked hydrogels led to almost uniformly collective strand migration from MCF10A spheroids, we next modulated bulk crosslinking/stiffness while maintaining a constant high density of fibers (3% vol). At low crosslinking/stiffness (1.5% LAP), predominantly single cell migration was observed as either amoeboid (red arrowheads) or elongated, single mesenchymal (yellow arrowheads) phenotypes (Fig. 5A). At an intermediate crosslinking/stiffness (2.0% LAP), increased heterogeneity in migration phenotypes was observed. Single cells migrating with rounded amoeboid or spread mesenchymal morphologies, collective strands, and disconnected multicellular clusters all emanated from the same spheroid. Each of these distinct migratory modes, except rounded amoeboid cells, appeared to directly engage fibers, as evidenced by proximity to and deformation of fibers (Fig. 5C, red arrowheads). In contrast to studies in nonfibrous GelMA (Fig. 3), we observed the highest total number of migrating cells at an intermediate gel stiffness (2.0% LAP) and equal total migration distance between low (1.5% LAP) and intermediate (2.0% LAP) gel stiffness (Fig. 5D-E). Interestingly, quantification of migration modes revealed that at any given gel stiffness the most prevalent migration mode corresponded to the greatest number of escaped cells and highest total migration distance (Fig. 5F). Single migrating cells were greatest in number and total migration distance at low stiffness/crosslinking (1.5% LAP), which favored single cell migration; similarly, these same metrics were greatest for multicellular clusters in intermediate stiffness gels (2.0% LAP) and for collective strands in high stiffness/crosslinking gels (2.5% and 3.0% LAP). This result suggests a propensity for migratory cells to optimize their migration mode to most effectively navigate a given microenvironment.

**Fig.5.**
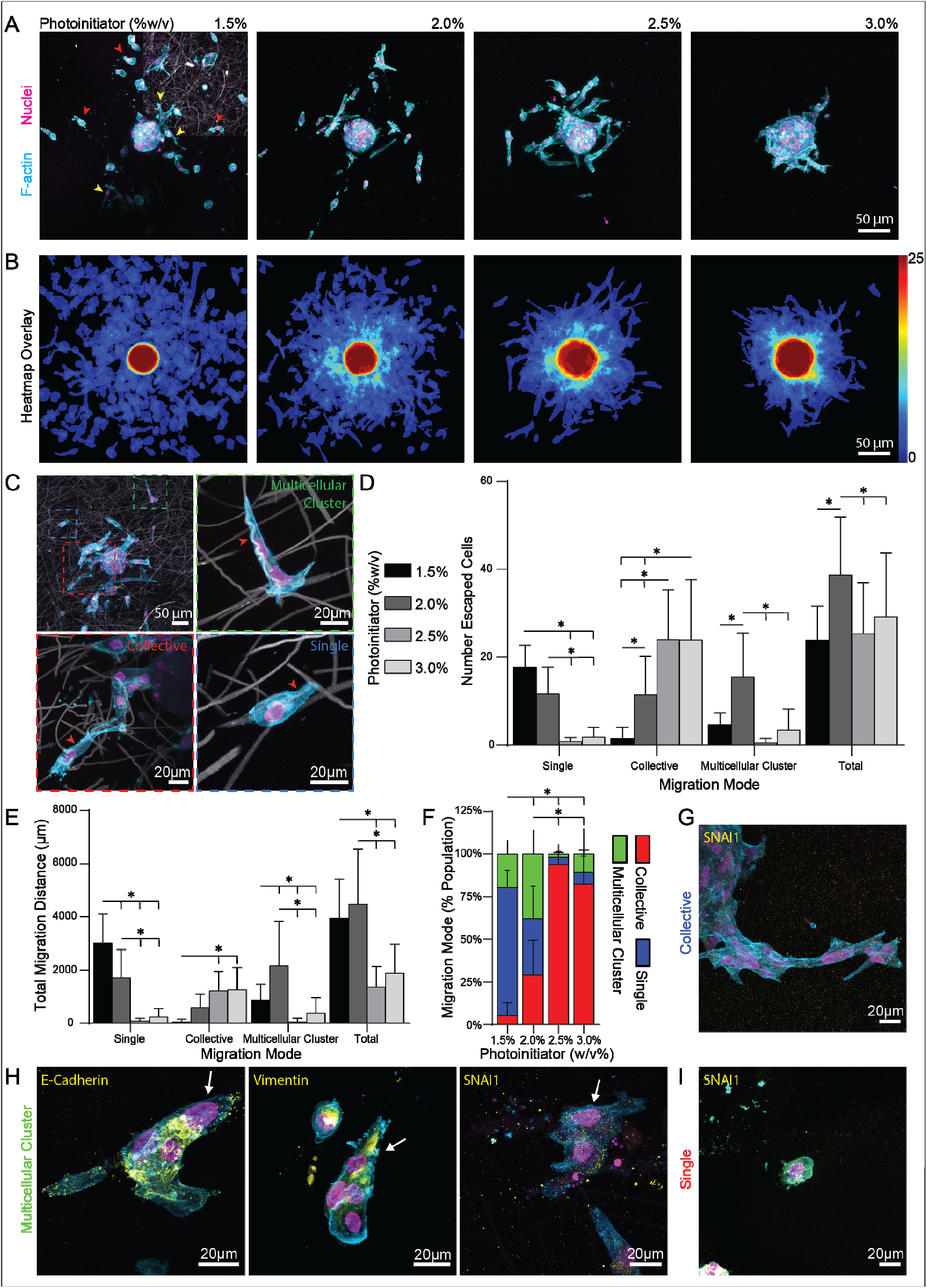
Decreased hydrogel stiffness with matrix fibers present elicits diverse migration modes. (A) Fluorescent images of spheroid outgrowth after 4 days in 3 vol% fiber density gels over a range of photoinitiator concentrations. (B) Corresponding heatmap overlays of binarized actin images for n = 25 spheroids per condition. (C) Higher magnification images of distinct migration modes engaging with DexVS fibers. Red arrowheads indicate fiber engagement. Quantification of (D) total number of escaped cells and (E) total migration distance per spheroid as a function of migration mode. (F) Stratification of migration mode subpopulations per spheroid. (G) Representative images of multicellular outgrowths immunostained for E-cadherin, vimentin, and SNAI1 in collective strands. Representative images of (H) multicellular clusters (white arrows indicated leading invasive edge) and (I) single cells immunostained for SNAI1. All data presented as mean ± std; * indicates a statistically significant comparison with p < 0.05.

At an intermediate level of crosslinking/stiffness (2.0% LAP), we observed the greatest degree of heterogeneity in migration mode, with single cells, collective strands, and multicellular clusters nearly equally represented across all invading cells. Multicellular clusters on average consisted of 2-4 nuclei with a prominent leading cell (white arrow) followed by several stalk cells (Fig. 5H, Supp. Fig. 2). EMT signaling has been strongly implicated in the regulation of cell-cell adhesion that could contribute to breakage events leading to the formation of multicellular clusters^22,23^. We immunostained collective strands and multicellular clusters for E-cadherin, vimentin, and SNAI1 (a key EMT transcription factor). Multicellular clusters expressed both E-cadherin at cell-cell junctions and fibrillar vimentin caging the nucleus, similar to collective strands (Fig. 4G & 5I). However, both collective strands and multicellular clusters expressed only low levels of cytosolic SNAI1 and no evidence of nuclear localization of this key EMT transcription factor (Fig. 5G-I), again suggesting that both migration phenotypes arise from incomplete or partial EMT.

### TGF-β1 elicits heterogeneous migration modes via EMT induction

The presence of fibrillar vimentin in collective strands and multicellular clusters suggests that these are EMT-driven migration modes, but likely involving only partial EMT given the maintenance of E-cadherin-mediated cell-cell connectivity (Fig. 4-5). To further examine whether EMT influences the migration modes observed in this model, we treated cultures with TGF-β1, a soluble factor known to drive EMT in MCF10As^24–26^. In fibrous, high crosslinking/stiffness gels that elicited predominantly collective strand migration in earlier studies (Fig. 4), TGF-β1 at any concentration increased the total number of escaped cells per spheroid (Fig. 6A-B). The majority of this increase resulted from significant increases in the number of cells migrating as collective strands and a trend towards an increase in the number of single cells (Fig. 6C). Interestingly, the maximum invasion depth of multicellular clusters proved insensitive to TGF-β1 concentrations while the two highest doses of TGF-β1 (2.0 and 10.0 ng ml^-1^) significantly increased single cell and collective strand invasion depth (Supp. Fig. 3). The two highest doses of TGF-β1 (2.0 and 10.0 ng ml^-1^) also led to increases in total migration distance, but only the highest dose (10 ng mL^-1^) led to a significant increase in migration distance in single cells (Fig. 6D). TGF-β1 at any concentration led to the emergence of a minor (2-5%) fraction of multicellular clusters. Overall, TGF-β1 increased the fraction of invading cells migrating as individuals with commensurate decreases in the frequency of collective strands (Fig. 6E). Interestingly, this transition is distinct to that observed in bulk stiffness-mediated collective to single mode switching, in which decreasing bulk stiffness of fibrous gels decreased collective strand migration but increased both multicellular cluster and single cell migration (Fig. 5H).

**Fig.6.**
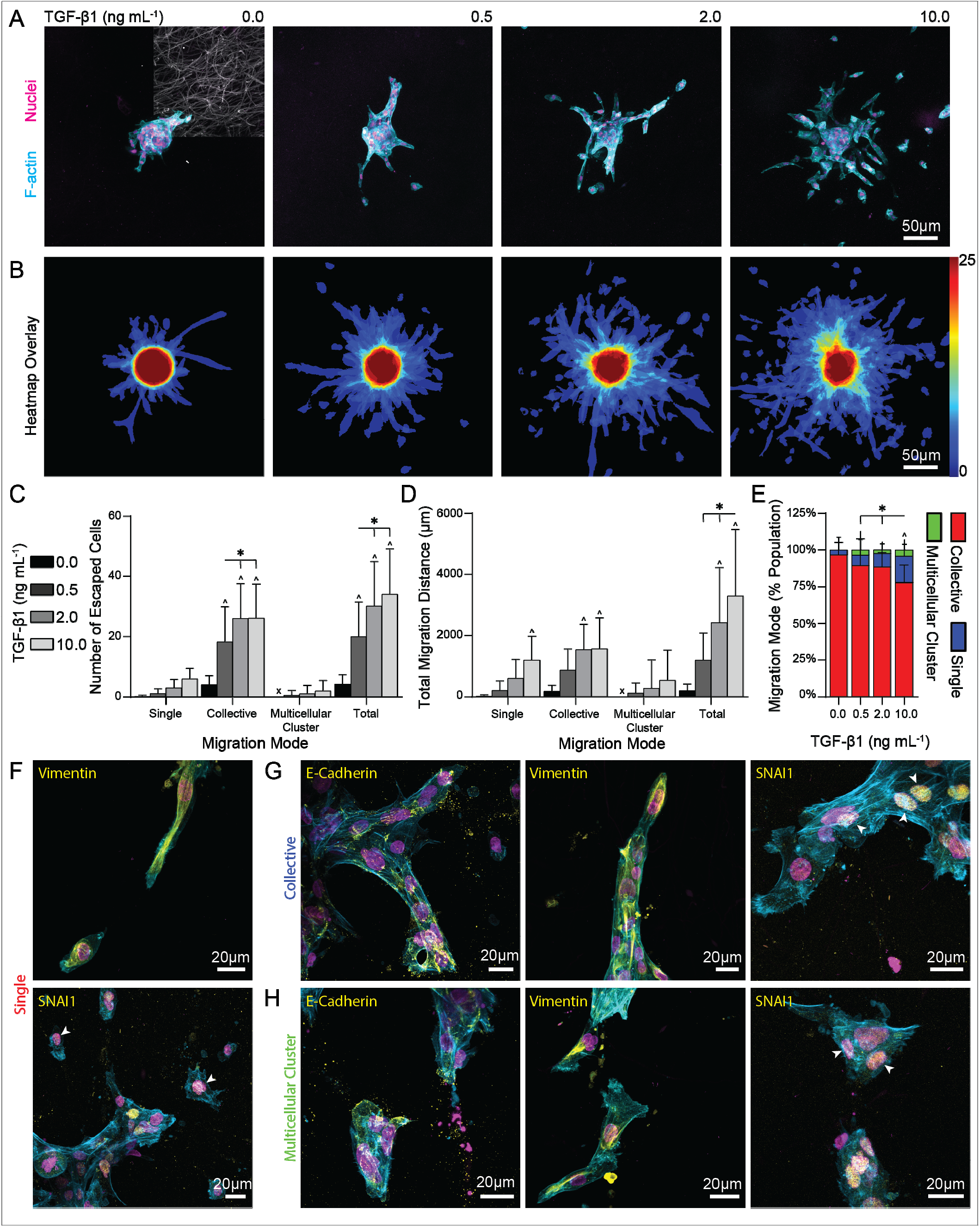
TGFB promotes MCF10A escape from spheroids and heterogeneity in migratory phenotypes. (A) Fluorescent images of 2 day spheroid outgrowth in stiff 2.5 % LAP, 3 vol% fiber density gels treated with soluble TGF-β1 in culture media. (B) Corresponding heatmap overlays of actin images for n = 25 spheroids per condition. Quantification of (C) total number of escaped cells and (D) total migration distance per spheroid as a function of migration mode. (E) Stratification of migration mode subpopulations per spheroid. EMT marker expression following 10 ng mL^-1^ TGF-β1 treatment in (F) single cell, (G) collective strand, and (H) multicellular cluster phenotypes. White arrowheads indicate SNAI1 nuclear expression. All data presented as mean ± std; * indicates a statistically significant comparison with p < 0.05; ^ indicates significance against 0 ng mL^-1^ TGF-β1.

Compared to changes in migration mode as a function of bulk hydrogel stiffness, TGF-β1-mediated migration suggested stronger involvement of EMT. Single cells, regardless of their proximity to a multicellular structure, possessed nuclear expression of SNAI1 (Fig. 6F, white arrowheads) in addition to fibrillar vimentin assembled around nuclei. Both collective strands and multicellular clusters expressed E-cadherin at cell-cell junctions, but with reduced levels compared to non-TGF-β1 controls. Both multicellular modes of migration again assembled vimentin networks at higher levels than non-migratory cells remaining within the spheroid (Supp. Fig. 4). Further supporting a more complete or active mesenchymal transition, nuclear localization of SNAI1 was observed in a subset of cells in all multicellular clusters (Fig. 6G-H, white arrowheads).

### Collective strands better resists apoptosis than single cells or multicellular clusters

A purported consequence of heterogeneity in cell migration mode is therapy resistance, where radio-or chemotherapy may prove efficacious in eliminating only a subset of migration modes^27–29^. Additionally, therapies may induce migration mode switching, diminishing the effectiveness of a given treatment as cells transition to a more resistant migratory phenotype^20^. To explore these possibilities, we leveraged our biomaterial system to assess apoptotic resistance as a function of migration mode. Single cells, collective strands, and multicellular clusters were produced in either fibrous, intermediate bulk hydrogel stiffness gels (Fig. 5) or fibrous, high bulk hydrogel stiffness gels cultured in 10 ng mL^-1^ TGF-β1 (Fig. 6). Following migration from spheroids, cultures were dosed with the established potent pro-apoptotic agent staurosporine^30^ for 6 hours and then assessed for cell death by caspase-3 immunostaining (Supp. Fig. 5A-B). In controls lacking staurosporine treatment, caspase-3 was observed at low baseline levels within nonmigratory cells of the spheroid body and migratory cells regardless of migration mode. In contrast, staurosporine-treatment significantly increased caspase-3 expression levels within the spheroid body and across all migration modes (Supp. Fig. 5C). In both gel conditions, staurosporine treatment had no effect on maximum invasion depth of collective strands (Supp. Fig. 5D) but drastically reduced the frequency of both single migrating cells and noncontiguous multicellular clusters (Fig. 7A-D). At a high staurosporine dose (10 nM) single cells and multicellular clusters were near completely abrogated (Fig. 7E, G), resulting in almost entirely collective strands (Fig. 7F, H). Morphologically, collective strands treated with staurosporine appeared thinner with disrupted actin networks and chromatin condensation, suggesting the onset of apoptosis but at insufficient levels to induce nuclear and cytoplasmic degradation (Fig. 7I). Interestingly, the number of cells migrating as collective strands remained unchanged, indicating preferential survival of collective strands over single cells or multicellular clusters following staurosporine treatment (Fig. 7E,G).

**Fig.7.**
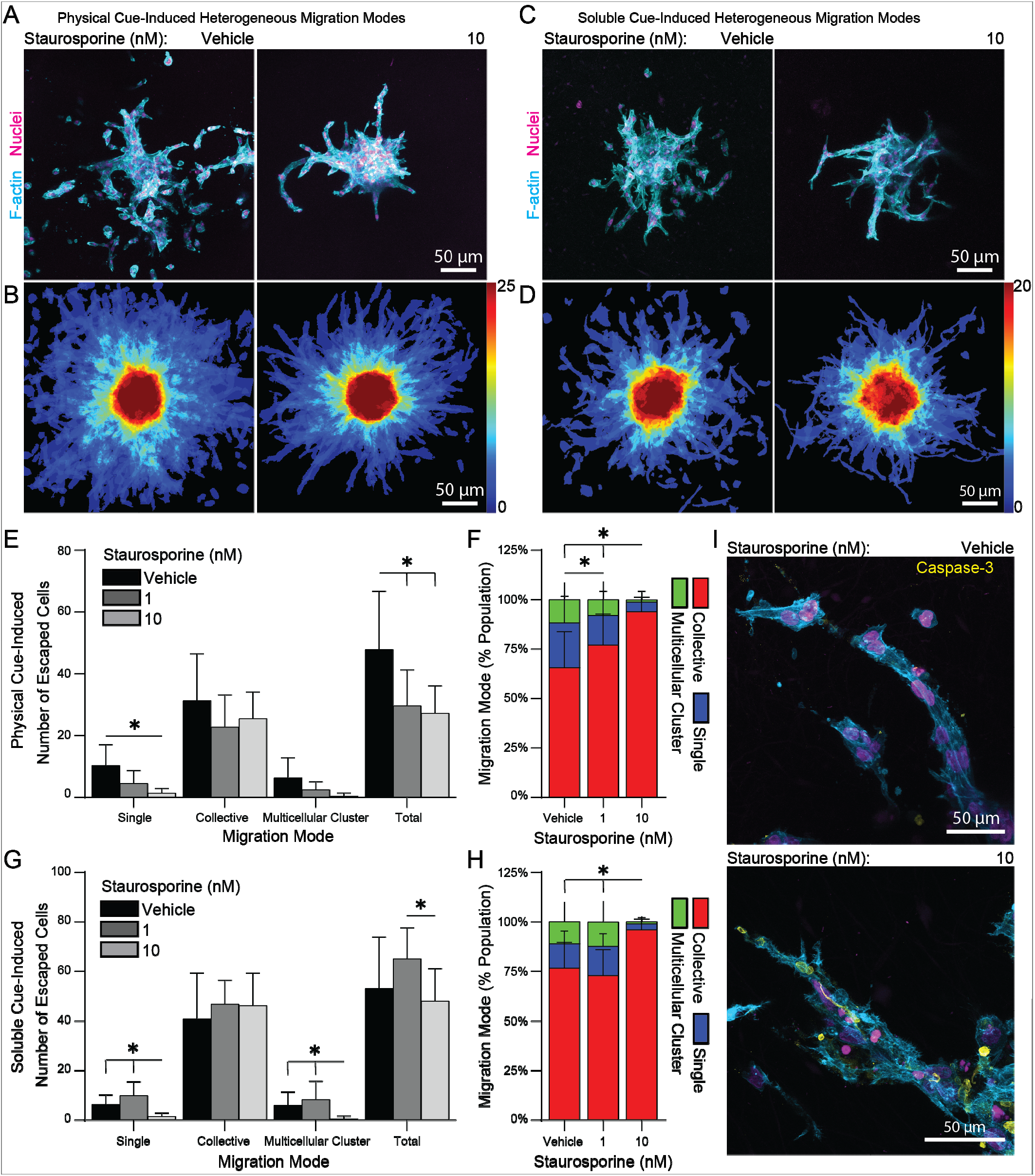
Cell migration mode defines differential susceptibility to apoptosis from staurosporine treatment. (A) Fluorescent images of spheroid outgrowth after 6 days in intermediate stiffness 2.0% LAP, 3 vol% fiber density gels and treated with staurosporine for 6 hours prior to fixation. (B) Corresponding heatmap overlays of binarized actin images for n = 25 spheroids per condition. (C) Fluorescent images of 4 day spheroid outgrowth in stiff 2.5% LAP, 3 vol% fiber density gels treated with 10 ng mL^-1^ TGF-β1 and pulsed with staurosporine for 6 hours prior to fixation. (D) Corresponding heatmap overlays of actin images for n = 20 spheroids per condition. Quantification of total number of escaped cells and migration mode subpopulations per spheroid for (E-F) physical-cue induced heterogeneous migration modes or (G-H) soluble-cue induced heterogeneous migration modes. (I) Caspase-3 expression and morphology of collective strands treated with 10 nM staurosporine. All data presented as mean ± std; * indicates a statistically significant comparison with p < 0.05.

## DISCUSSION

Here, we describe a composite biomaterial consisting of synthetic DexVS fibers embedded in amorphous GelMA hydrogel to generate biomimetic stromal matrices in which biophysical and biochemical cues relevant to cancer can be orthogonally tuned. This biomaterial system enabled new insights into how matrix stiffness and fiber density, two physical cues that vary with tumor progression, individually influence invasive potential, EMT, and migration phenotypes of otherwise quiescent MCF10As. We found that bulk hydrogel stiffness/crosslinking and fiber density distinctly influence migration modes that have been commonly reported *in vivo* including amoeboid, single mesenchymal, multicellular clusters, and collective strands^29,31^. Further, we identified microenvironmental conditions that elicit controlled subpopulations of these various migration strategies. While previous studies attribute distinct migration modes to cell-specific genetic variation through the differential accrual of mutations^32^, this work supports the notion that physical cues from the matrix can define heterogeneity in cell migration mode despite an initially homogeneous cell population.

We first modulated the degree of crosslinking of GelMA hydrogels lacking fibrous architecture to determine the influence of hydrogel stiffness on cell migration. Despite the presence of endogenous cell-adhesive RGD within GelMA^33^, we observed solely amoeboid migration, which increased in frequency with decreasing GelMA crosslinking/stiffness (Fig. 3). Timelapse imaging confirmed cells underwent cyclic blebbing and rapid squeezing events to migrate, perhaps due to the nanoporosity of GelMA hydrogels leading to cell confinement^6,34^. While migration in nonfibrous GelMA was primarily amoeboid, we observed a shift towards mesenchymal cell migration with the inclusion of cell-adhesive matrix fibers (Fig. 5). This transition may result from enhanced cell adhesion to more rigid RGD-containing fiber segments^35^.

The DexVS fibers employed in this 3D stromal mimetic were designed to model fibrous collagens found within the peritumoral stroma, which have been widely implicated in cancer cell migration via contact guidance^36^. DexVS fibers were functionalized with RGD given previous implications that RGD-binding integrins are required for cancer cell migration^37^. Additionally, RGD is the predominant adhesive ligand present in gelatin and GelMA hydrogels^33^, therefore allowing us to maintain a consistent adhesive ligand across bulk hydrogel and fibrous components. Thus, cells could interact with both structural components without altering expression of integrin subtypes. DexVS fibers can be readily functionalized with other thiolated peptides via Michael-type addition, including GxOGER to model stromal type I collagen and other full-length proteins^38^. Future work incorporating other integrin-binding ligands in a controlled manner could enable systematic investigation into the influence of ECM composition to cancer progression.

To our surprise, cell adhesion to matrix fibers was a requirement for multicellular migration strategies from MCF10A spheroids (Fig. 4). The observed coordination of multicellular invasion may arise from fiber-mediated contact guidance that promotes directional streaming and/or enhanced contractility that acts to maintain the integrity of cell-cell adhesions^39,40^. Modulating gel stiffness while maintaining a constant fiber density engendered migration heterogeneity including amoeboid, single mesenchymal, multicellular clusters, and collective strands (Fig. 5). Whether an expanded repertoire of distinct migration modes contributes greater metastatic burden compared to a more homogeneous tumor is an important outstanding question. Previous *in vivo* studies suggest migration mode can select for hematogenous vs. lymphatic dissemination^41^ and that intravasation efficiency is a function of migration mode^42^. The integration of the material system presented here with organotypic or microfluidic platforms that model intravasation events could more directly address this question.

EMT is widely implicated in metastasis and has recently been described as a continuum (rather than a binary transition) with step-wise programming events that cumulatively contribute to invasive behavior^43,44^. As such, we sought to characterize the degree of EMT across the different modes of migration observed in our model. Multicellular migration modes in fibrous matrices retained epithelial marker E-cadherin while adopting mesenchymal maker vimentin (Fig. 4-5), suggesting that adhesion to matrix fibers promotes at least partial EMT programming. While a significant prior body of work has focused on soluble factors driving EMT in cancer (eg. TGF-β1, EGF, FGF), more recent evidence indicates that physical cues from the matrix can also potentiate EMT^17,26,45^. Whether fibers promote EMT and migration through topographical effects and/or by providing stiff microdomains that activate mechanosensing pathways remains to be explored^46,47^. To fully promote a mesenchymal state and establish an EMT migration mode in our model, we subjected MCF10A spheroids to TGF-β1. We observed a dose-dependent increase in cell migration and the emergence of diverse migration modes (Fig. 6). Interestingly, nuclear localization of SNAI1 was comparable across all invasive structures and not solely evident in single migrating cells. It is possible that SNAI1 primes cells within collective strands to transition to single cell migration through decreasing connectivity between cells, evident in decreased E-cadherin in collective strands following TGF-β1 treatment. Understanding how stromal biophysical cues potentiate pro-metastatic cytokines to drive EMT and invasive behavior could help inform cancer therapies targeting physical matrix cues in the tumor microenvironment^48^.

A purported functional consequence of migration mode is resistance to radio- and chemotherapies^27–29,49^. Here, we observed that collective strands were more resistant to staurosporine-induced apoptosis than either single cells or multicellular clusters (Fig. 7). Migration mode defines adhesion-mediated survival pathways which may contribute to therapy resistance^21^. Recently, Haeger et al. demonstrated a similar trend in murine orthotopic sarcoma and melanoma models in which collective strands better resisted radiation therapy and DNA damage compared to single cells^27^. Plasticity following therapy may also contribute to apoptosis resistance. Wolf et al. and others have demonstrated anti-MMP therapeutics targeting single mesenchymal invasion can trigger mesenchymal-to-amoeboid transition, enabling therapy evasion^18,50^. Overcoming these plasticity-based therapy resistances will likely require an enhanced understanding of molecular differences between each mode of migration. Control over a range of migration modes within our stromal mimetic enables systematic investigation of EMT-TFs, cell-cell and cell-ECM adhesion molecules, and mechanotransduction proteins like YAP/TAZ that could underlie migration mode heterogeneity.

## CONCLUSION

We present an *in vitro* stromal mimetic in which biomechanical and biochemical cues can be orthogonally modulated to elicit a range of migration modes relevant to cancer cell transtromal migration. We show how two key features, fiber density and bulk stiffness, independently and in combination affect cell migratory capacity and dictate mode switching events between amoeboid, single mesenchymal, multicellular clusters, and collective strands. We find that cell adhesion to ECM fibers induce a partial-EMT state, resulting in an invasive multicellular strand-like migration mode, while further promoting a mesenchymal phenotype via a soluble cue drives single cell migration. Furthermore, we show that migration as collective strands provides enhanced protection against a potent apoptotic signal as compared to single cells and multicellular clusters. This model could help elucidate molecular pathways that govern changes in cell migration modes and resulting heterogeneity.

## ACKNOWLEDGEMENTS

DLM, WYW, and JMB acknowledge financial support from the National Science Foundation Graduate Research Fellowship Program (DGE1256260). W.Y.W acknowledges financial support from the University of Michigan Rackham Merit Fellowship. B.M.B. acknowledges financial support from an NIH Pathway to Independence Award (HL124322). The authors acknowledge funding from United States National Institutes of Health grants R01CA238042, R01CA196018, U01CA210152, R01CA238023, R33CA225549, R50CA221807, and R37CA222563. Brock Humphries, PhD, was supported by an American Cancer Society -Michigan Cancer Research Fund Postdoctoral Fellowship, PF-18-236-01-CCG.

## DATA AVAILABILITY

The data that support the findings of this study are available from the corresponding author upon reasonable request.

## Notes

### Competing Interest Statement

The authors have declared no competing interest.

