## Supplemental Figures for "Fiber density and matrix stiffness modulate distinct cell migration modes in a 3D stroma mimetic composite hydrogel"

**Supplementary Material**

The Supplementary Material includes 5 Supplementary Figures and 1 Supplementary Video

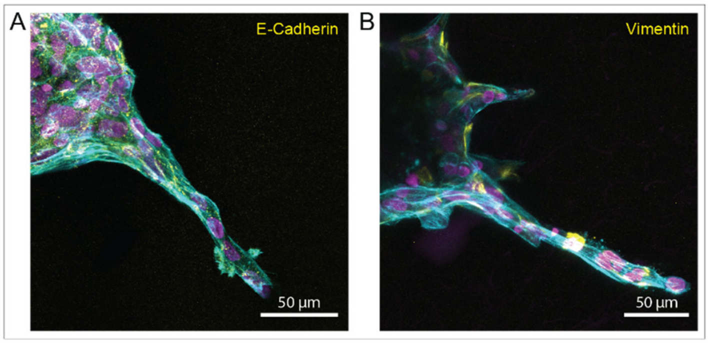


**Supp. Fig. 1. Fluorescent images of spheroid outgrowth over 4 days in stiff 2.5% LAP, 3 vol% fiber density gels. EMT marker expression in collective strands and the spheroids body showing (A) E-cadherin and (B) vimentin.**

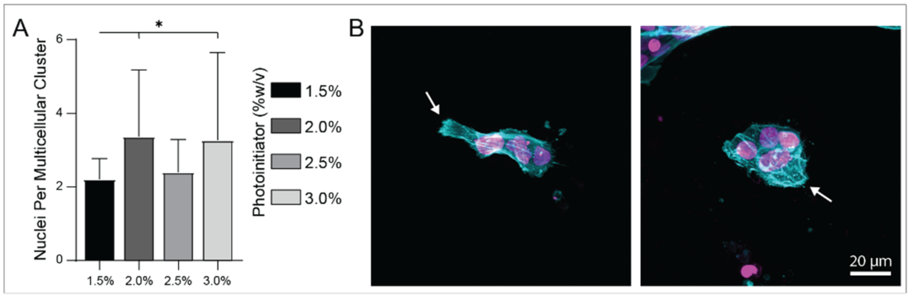


**Supp. Fig. 2. (A) Quantification of nuclei per multicellular cluster outgrowth in 3 vol% fiber density gels over a range of photoinitiator concentrations. (B) Fluorescent images of**

**distinct multicellular cluster morphologies. White arrows indicate leading invasive edge of multicellular clusters.**

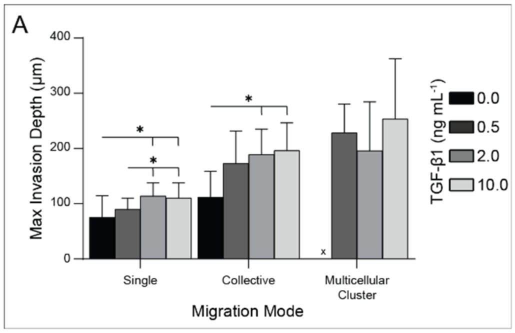


**Supp. Fig. 3. (A) Quantification of maximum invasion depth of distinct migration modes in stiff 2.5% LAP, 3 vol% fiber density gels treated with soluble TGF-β1 in culture media.**

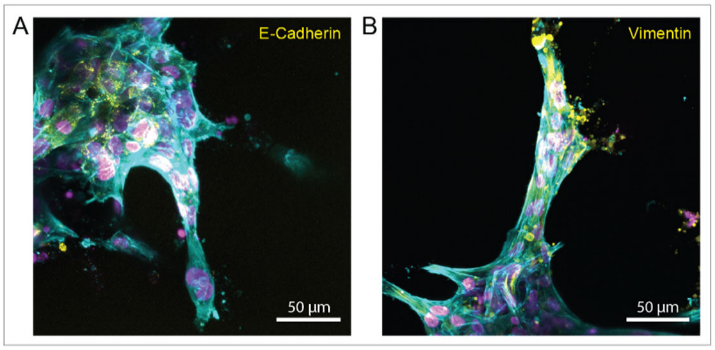


**Supp. Fig. 4. Fluorescent images of spheroid outgrowth over 4 days in stiff 2.5% LAP, 3 vol% fiber density gels treated with 2 ng mL$^{-1}$ TGF-β1 in culture media. EMT marker expression in collective strands and the spheroids body showing (A) E-cadherin and (B) vimentin.**

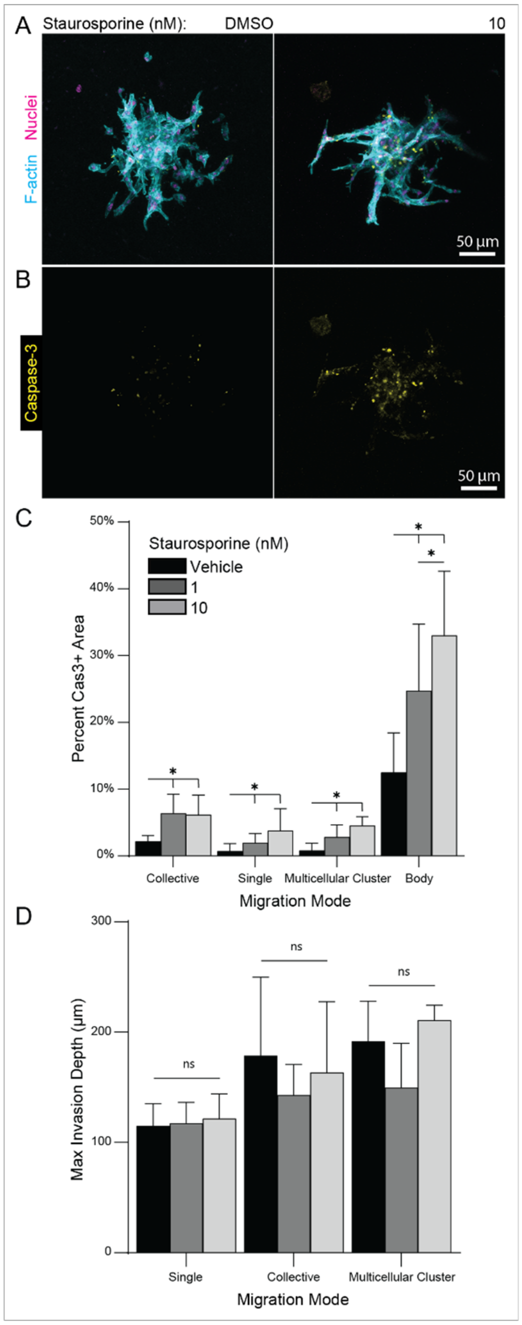


**Supp. Fig. 5. Quantification of (A) caspase-3 expression and (B) maximum invasion depth across distinct migration modes in stiff 2.5% LAP, 3 vol% fiber density gels pulsed with staurosporine for 6 hours prior to fixation.**

**Supp. Video 1. Time-lapse images of LifeAct-GFP expressing MCF10A cells migrating in nonfibrous low stiffness (1.5% LAP) gels.**
